# Acetylcholine synergizes with netrin-1 to drive persistent firing in the entorhinal cortex

**DOI:** 10.1101/2020.07.08.194274

**Authors:** Stephen D. Glasgow, Edwin W. Wong, Ian V. Beamish, Kevin Lancon, Julien Gibon, Philippe Séguéla, Edward S. Ruthazer, Timothy E. Kennedy

**Affiliations:** Department of Neurology & Neurosurgery, Montreal Neurological Institute-Hospital, McGill University, Montréal, QC, H3A 2B4, Canada; Department of Biology, University of British Columbia, Kelowna, BC, V1V 1V7, Canada

## Abstract

The ability of the mammalian brain to maintain spatial representations of external or internal information for short periods of time has been associated with sustained neuronal spiking and reverberatory neural network activity in the medial entorhinal cortex. Here, we show that cholinergic activation of muscarinic receptors on entorhinal cortical neurons mediates plasma membrane recruitment of the netrin-1 receptor deleted-in-colorectal cancer (DCC) to promote muscarinic receptor-mediated persistent firing. Conditional deletion of netrin-1 or DCC, which are required for synaptic plasticity, inhibits cholinergic persistent firing, and leads to deficits in spatial working memory. Together, these findings indicate that normal working memory function requires synergistic action of acetylcholine and netrin-1.

---

The entorhinal cortex is a critical node within the hippocampal formation, and serves as a functional hub to a widespread network of circuits involved in spatial memory (Eichenbaum, 2000). Periods of sustained neural activity in the entorhinal cortex have been reported during working memory tasks, suggesting neuronal firing may form the cellular basis of certain forms of short-term memory (Bittner et al., 2017; Hasselmo, 2008). Layer V entorhinal neurons are endowed with the ability to generate persistent neuronal firing activity, a form of intrinsic cellular plasticity characterized by plateau potentials and long-lasting periods of cellular spiking that greatly outlast initial stimulation (Egorov et al., 2002). This persistent firing is triggered by activation of muscarinic cholinergic receptors, leading to increased levels of intracellular Ca^2+^ ([Ca^2+^]_i_) (Haj-Dahmane and Andrade, 1999; Ratte et al., 2018; Zhang et al., 2011).

Netrin-1 binds to its canonical receptor, deleted-in-colorectal cancer (DCC), to activate phospholipase C (PLCγ), trigger release of [Ca^2+^]_i_ through inositol triphosphate receptors (IP_3_Rs), and activate canonical transient receptor potential channel 1 (TRPC1) (Kang et al., 2018; Venkatachalam et al., 2003; Wang and Poo, 2005). Netrin-1 is secreted from dendrites in response to neuronal activation and neuronal expression of netrin-1 is required for activity dependent long-term potentiation (LTP) (Glasgow et al., 2018). Netrin-1 and DCC are expressed throughout all layers of the adult entorhinal cortex (Lein et al., 2007), suggesting that netrin-1 may contribute to gating cellular excitability in the entorhinal cortex during working memory by promoting persistent firing activity.

To assess whether netrin-1 impacts membrane depolarization in the mature mammalian nervous system, we initially performed whole-cell current-clamp recordings from layer V entorhinal neurons from adult mice in the presence of kynurenic acid (KYNA; 1 mM) and picrotoxin (PTX; 100 μM) to block fast ionotropic glutamatergic and GABAergic synaptic transmission, respectively. Bath application of netrin-1 (2.7 nM) resulted in a significant, dose-dependent depolarization of membrane potential (Fig. 1A-B). Interestingly, bath application of netrin-1 did not result in significant changes in input resistance, but resulted in minor alterations in other electrophysiological properties associated with changes in *I*_h_, a voltage-dependent mixed cationic current that contributes to persistent firing and working memory function in the prefrontal cortex (Fig. 1C-F) (Thuault et al., 2013). However, netrin-1 failed to induce any other detectable changes in *I*_h_, indicating that alteration of *I*_h_ is unlikely to be responsible for netrin-1 depolarization of membrane potential (Fig. S1).

**Fig. 1.**
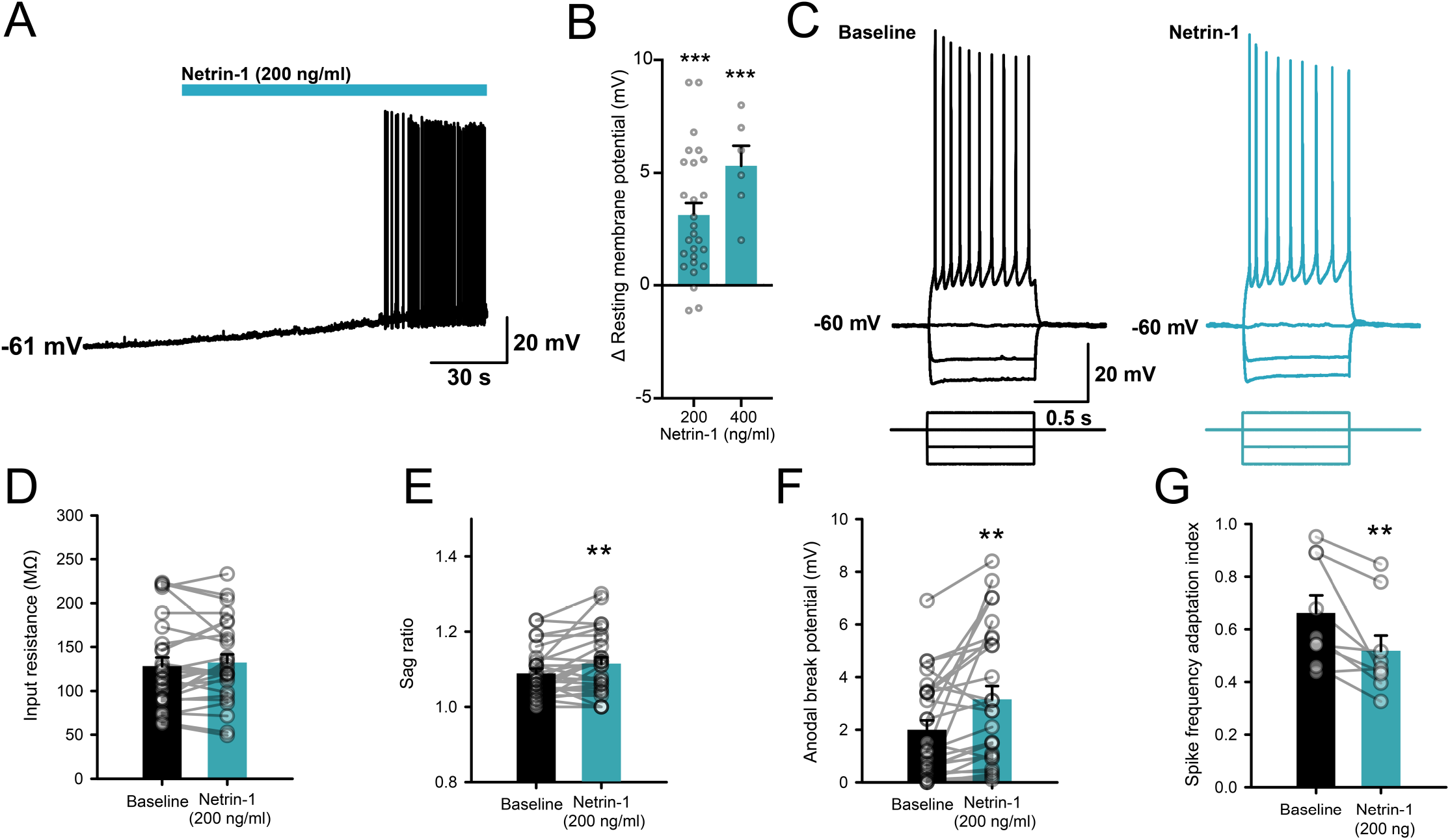
Netrin-1 depolarizes layer V entorhinal neurons. A. Membrane potential recording showing that bath application of netrin-1 (2.7 nM, 2 min) results in a long-lasting depolarization in layer V entorhinal neurons. B. Group data showing that netrin-1 results in a dose-dependent depolarization of membrane potential (2.7 nM: 3.1±0.9 mV, 5.3 nM: 5.3±0.9 mV). C. Membrane response to a series of hyperpolarizing and depolarizing current steps in a layer V entorhinal pyramidal neuron prior to (left, black) and following bath application of netrin-1 (2.7 nM; right, blue). D-G. Group data showing the effect of netrin-1 (2.7 nM) on input resistance (D), sag ratio (E; 1.09±0.01 in control vs. 1.12±0.02 in netrin-1), anodal break potential (F; 1.93±0.37 mV in control vs. 3.15±0.50 in netrin-1), and spike frequency adaptation index (G; 0.66±0.07 in control vs. 0.52±0.06 in netrin-1). * denotes *p*<0.05, ** denotes *p*<0.01, *** denotes *p*<0.001.

In developing axons, increases in [Ca^2+^]_i_ levels through L-type Ca^2+^ and TRPC channels are critical mediators of netrin-1 depolarization of the growth cone (Magistretti et al., 2004; Wang and Poo, 2005). To test if netrin-1 increases [Ca^2+^]_i_ in cortical neurons, we recorded somatic [Ca^2+^]_i_ levels in dissociated cortical neurons following application of netrin-1. Brief bath application of netrin-1 (2.7 nM) resulted in a rapid and long-lasting increase in [Ca^2+^]_i_ in cultured cortical neurons. This effect was dependent on extracellular Ca^2+^, as well as partially-dependent on L-type Ca^2+^ channels and an additional non-specific cationic channel, as well as being mediated at later timepoints through modulation of [Ca^2+^]_i_ stores (Fig. S2). Together, these findings suggest that regulation of [Ca^2+^]_i_ contributes to the expression of netrin-1-mediated inward currents.

To characterize netrin-1-mediated membrane currents, we performed whole-cell voltage clamp recordings from layer V neurons in acute entorhinal brain slices from adult mice. Netrin-1 induced a long-lasting inward current at a resting potential of −70 mV. The current-voltage relationship was biphasic and reversed near −41 mV (goodness of fit for Boltzmann sigmoidal, *R^2^*=0.494), consistent with activation of a calcium-activated, non-selective cationic channel (*I*_CAN_) (Fig. S3) (Congar et al., 1997). Interestingly, blockade of L-type Ca^2+^ channels using nifedipine (10 μM) failed to inhibit the netrin-1 current, whereas conditional deletion of the netrin-1 receptor DCC as well as pharmacological blockade of cationic channels, including TRPC1, with SKF96365 (100 μM) significantly attenuated the netrin-1 current (Fig. S3). These findings suggest that DCC-mediated increases in [Ca^2+^]_i_ activate currents that underlie netrin-1 depolarization of layer V entorhinal neurons.

Increased [Ca^2+^]_i_ through activation of calcium-activated non-selective cation current (*I*_CAN_) is known to contribute to persistent firing of cortical neurons, which has been proposed as a cellular basis of working memory (Thuault et al., 2013; Zhang et al., 2011). Our findings suggest that netrin-1 alone might be sufficient to promote persistent firing activity. To test this directly, we recorded from layer V entorhinal neurons, and attempted to induce persistent firing prior to and following bath application of netrin-1 (2.7 nM) in the presence of synaptic blockers. Interestingly, bath application of netrin-1 induced persistent firing in a small subset of layer V entorhinal neurons (1/11, 9.0%). In contrast, bath application of netrin-1 routinely led to depolarization of membrane potential and enhanced the amplitude of the slow afterdepolarization (sADP) that has been associated with *I*_CAN_-mediated emergence of plateau potentials and persistent firing (Fig. 2A-D) (Egorov et al., 2002; Zylberberg and Strowbridge, 2017).

**Figure 2.**
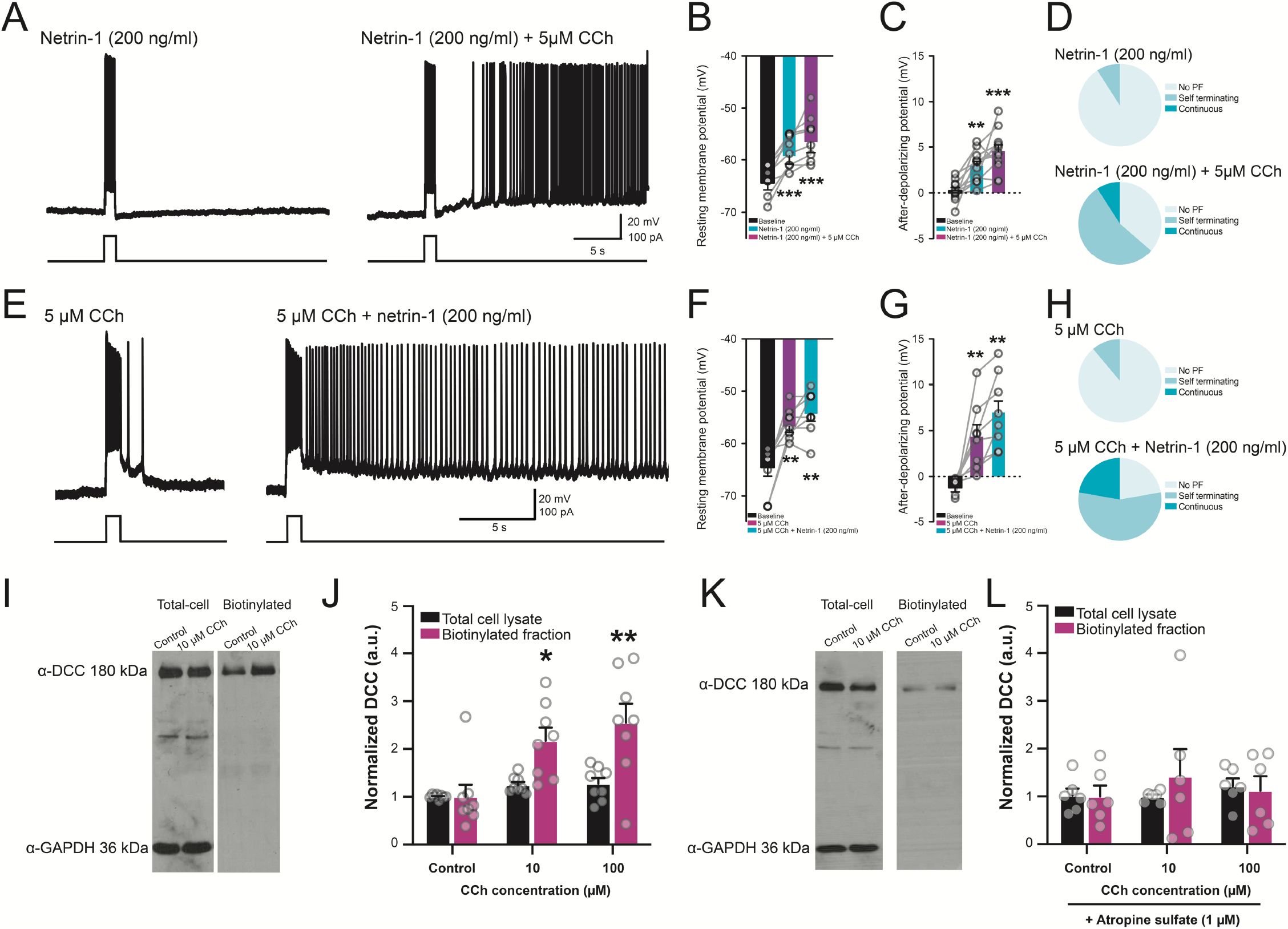
Netrin-1 potentiates cholinergically-mediated persistent firing in layer V entorhinal neurons. A. Bath application of netrin-1 (2.7 nM) alone fails to promote persistent firing activity (left), but co-application with CCh (5 μM, right) elicits plateau potentials and repetitive spiking activity. B-D. Group data show that netrin-1 and co-application with CCh (5 μM) increases resting membrane potential (B) and afterdepolarization potential (C; 0.22±0.33 mV in control, 2.91±0.51 mV in netrin-1, 4.51±0.70 mV in netrin-1 and 5 μM CCh), and induces persistent firing activity following +100 pA intracellular current injection in a large proportion of cells tested (D). E. Bath application of low concentrations of CCh (5 μM, left) fails to elicit persistent firing in response to a 1-s +100 pA depolarizing intracellular current pulse (left) but further co-application of netrin-1 (2.7 nM) results in the induction of a plateau potential and repetitive spiking (right). F-H. Group data show that CCh (5 μM) and co-application with CCh (5 μM) increases resting membrane potential (F; −64.6±1.7 mV in control, −56.8±1.1 mV in 5 μM CCh, −53.7±1.2 mV in 5 μM CCh and netrin-1), afterdepolarization potential (G; (−1.23±0.46 mV in control, 4.32±1.31 mV in 5 μM CCh, 6.97±1.26 mV in 5 μM CCh and netrin-1), and induces persistent firing activity following +100 pA intracellular current injection in the majority of cells tested (H; 1/9 cells, 11.1% in 5 μM CCh vs. 7/9 cells, 77.8% in 5 μM CCh and netrin-1). I. Representative Western blots (I) and group data (J) show that bath application of CCh results in a dose-dependent increase in surface distribution of DCC (blue bars) with no change in total DCC protein (black bars). K. Representative Western blots (K) and group data (L) of total (left in K, black bars in L) and surface biotinylated (right in K, blue bars in L) protein following bath application of CCh (0.1-100 μM) in the presence of muscarinic receptor antagonist atropine sulfate (1 μM). * denotes *p*<0.05, ** denotes *p*<0.01, *** denotes *p*<0.001.

High concentrations of the cholinergic agonist, carbachol (CCh; >10 μM), lead to a potent enhancement of sADP that is mediated, in part, through PLC-dependent actions on TRPCs (Egorov et al., 2002; Reboreda et al., 2018; Yan et al., 2009). However, lower concentrations of CCh more accurately reflect physiological levels of cholinergic receptor activation (Galvin et al., 2020; Marrosu et al., 1995). We postulated that netrin-1 may facilitate persistent firing activity in the presence of low concentrations of CCh (5 μM). Indeed, CCh (5 μM) together with netrin-1 further increased sADP amplitude (Fig. 2D). Importantly, co-application of CCh (5 μM) and netrin-1 (2.7 nM) routinely elicited long-lasting persistent firing activity in layer V entorhinal neurons (7/11 cells, 63.6%). Together, these data suggest that netrin-1 signaling can facilitate cholinergic neuromodulation to promote sustained neural activity in layer V entorhinal neurons.

To test whether lower levels of cholinergic receptor activation alone can enhance sADP amplitude and persistent firing activity, we applied 5 μM CCh in the presence of synaptic blockers. CCh (5 μM) led to membrane depolarization and an increase in sADP but failed to elicit any form of persistent firing in the majority of neurons recorded (8/9 cells, 89.9%). However, consistent with a synergistic relationship with cholinergic signaling, co-application of netrin-1 (2.7 nM) with CCh (5 μM) enhanced the sADP, and routinely triggered long-lasting plateau potentials and repetitive spiking activity (7/9 cells, 77.8%) (Fig. 2E-H). These findings indicate that cholinergic receptor activation and netrin-1 signaling combine to enhance sADP that can trigger long-lasting periods of depolarization and spiking in layer V entorhinal neurons.

Membrane potential depolarization potently regulates cell surface distribution of DCC, suggesting that cholinergic receptor activation may promote DCC trafficking to sensitize entorhinal neurons to netrin-1 (Bouchard et al., 2008). To determine whether cholinergic receptor activation recruits DCC to the plasma membrane, we used cell surface biotinylation assays for DCC following bath application of CCh. Brief cholinergic receptor activation (0.1-100 μM for 15 min) significantly increased plasma membrane DCC (Fig. 2I-J). The observed increase in surface DCC was blocked through pre-exposure to the muscarinic receptor antagonist, atropine sulfate (1 μM), indicating that this effect is mediated through activation of muscarinic receptors (Fig. 2K-L). Together, these findings suggest that cholinergic neurotransmission facilitates netrin-1 function by trafficking DCC to the cell surface.

To test whether the relationship between netrin-1 and cholinergic receptor activation was mediated through DCC, we conditionally-deleted DCC from principal excitatory forebrain neurons using a CaMKIIα-Cre driver line (CaMKIIα-Cre/DCC^*fl/fl*^; DCC cKO). We detected no differences in basal excitability between layer V entorhinal neurons from control and DCC cKO mice (Fig. 3A-G). Bath application of CCh (5 μM) resulted in significant depolarization of resting membrane potential and enhanced sADP in neurons from both control and DCC cKO mice, indicating that typical cholinergic receptor signaling remained intact in animals lacking DCC. However, we found that further co-application of netrin-1 enhanced sADP and drove depolarization in wildtype neurons but not in neurons from DCC cKO (Fig. 3H-J). Moreover, while co-application of 5 μM CCh and netrin-1 routinely induced persistent firing in layer V entorhinal neurons from wildtype control mice (8/12 cells, 67%), this same cocktail failed to induce persistent firing in any neurons from DCC cKO (0/12 cells, 0%). In contrast, increasing CCh concentration (10 μM) was sufficient to induce persistent firing in both wildtype control and DCC cKO entorhinal neurons (WT: 8/12 cells, 67%; DCC cKO: 9/11 cells, 75.0%) (Fig. S4). These findings indicate that DCC signaling contributes to the emergence of persistent firing activity in layer V entorhinal neurons, but that it is still possible to drive sustained firing and plateau potentials using high concentrations of cholinergic receptor agonist.

**Figure 3.**
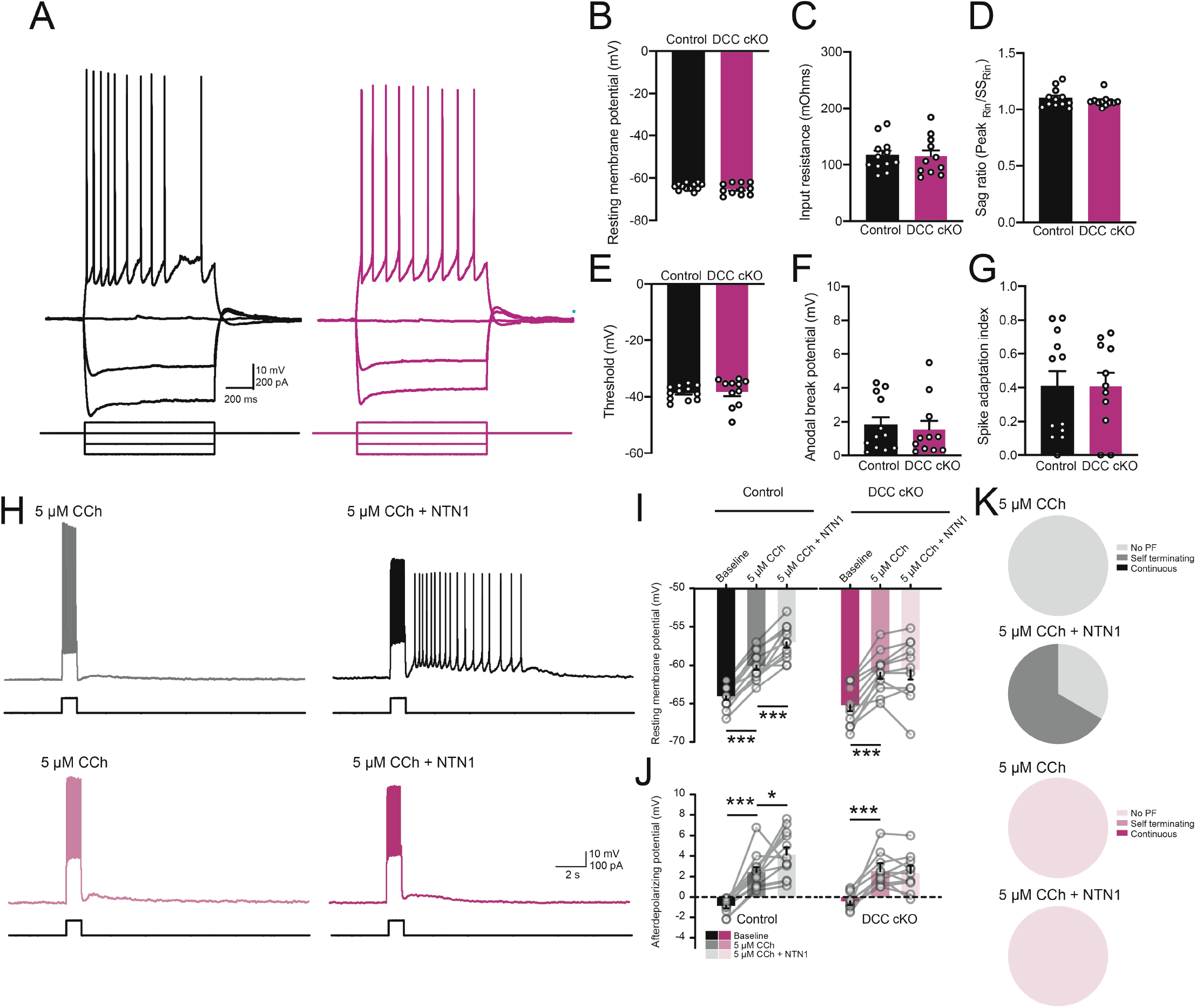
DCC is required for netrin-1 sensitization of CCh-induced persistent firing. A. Membrane voltage response to a series of depolarizing and hyperpolarizing current pulses (1 s) in layer V entorhinal neurons from control (black) or DCC cKO (purple) mice. B-G. Group data show no significant differences in resting membrane potential (B), input resistance (C), sag ratio (D), spike threshold (E), anodal break potential (F), or spike frequency adaptation index (G) between control (black) and DCC cKO (purple). H. Co-application of low concentrations of CCh (5 μM) and netrin-1 (2.7 nM) results in persistent firing (right) in control littermates (black, top) but not in layer V entorhinal neurons from DCC cKO (purple, bottom). I-K. Group data show that bath application of CCh (5 μM, red) and netrin-1 (2.7 nM, blue) results in significant membrane depolarization (I; WT: −64.1±0.5 mV in baseline, 60.0±0.5 mV in 5 μM CCh, 57.0±0.7 mV in 5 μM CCh and netrin-1; DCC cKO: −65.2±0.8 in baseline, 60.9±0.8 mV in 5 μM CCh, −60.9±0.8 mV in 5 μM CCh and netrin-1) and afterdepolarizing potential (J; WT: −0.87±0.21 mV in baseline, 2.42±0.48 mV in 5 μM CCh,4.11±0.70 mV in 5 μM CCh and netrin-1; DCC cKO: −0.41±0.36 in baseline, 2.53±0.77 mV in 5 μM CCh, 2.34±0.69 mV in 5 μM CCh and netrin-1). This triggered persistent firing activity (K) in control littermates (black, 8/12 cells, 66.7%) but not DCC cKO (purple, 0/11 cells, 0%). * denotes *p*<0.05, ** denotes *p*<0.01, *** denotes *p*<0.001.

DCC is activated by a number of ligands, including netrins, draxins, and cerebellins, suggesting other proteins may contribute to DCC-dependent persistent firing (Ahmed et al., 2011; Haddick et al., 2014; Wei et al., 2012). To examine the role of endogenous netrin-1 signaling on cholinergic receptor-mediated sustained spiking, we tested persistent firing activity in layer V entorhinal neurons conditionally-lacking netrin-1 expression by forebrain principal excitatory neurons (CaMKIIα-Cre/NTN1^*fl/fl*^ NTN1 cKO) (Glasgow et al., 2018). No observable changes in intrinsic cellular properties were noted between layer V neurons from either control or NTN1 cKO mice under baseline conditions (Fig. 4A-G). To determine whether netrin-1 contributes to cholinergic persistent firing activity, we bath applied high concentrations of CCh (10 μM) to layer V entorhinal neurons. Bath application of CCh (10 μM) led to a depolarization of membrane potential in both wildtype and NTN1 cKO neurons. In contrast, CCh enhanced sADP amplitude in layer V entorhinal neurons from wildtype mice but not NTN1 cKO mice (Fig. 4H-J). Moreover, CCh elicited persistent firing and plateau potentials in age-matched controls in the majority of neurons from wildtype mice (11/16 cells, 68.8%), but failed to induce spiking in most neurons from NTN1 cKO mice (PF triggered in 2/16 cells, 11.1%) (Fig. 4K). Together, these data point to a critical role for netrin-1 in the expression of persistent firing activity, which has been implicated in working memory function.

**Figure 4.**
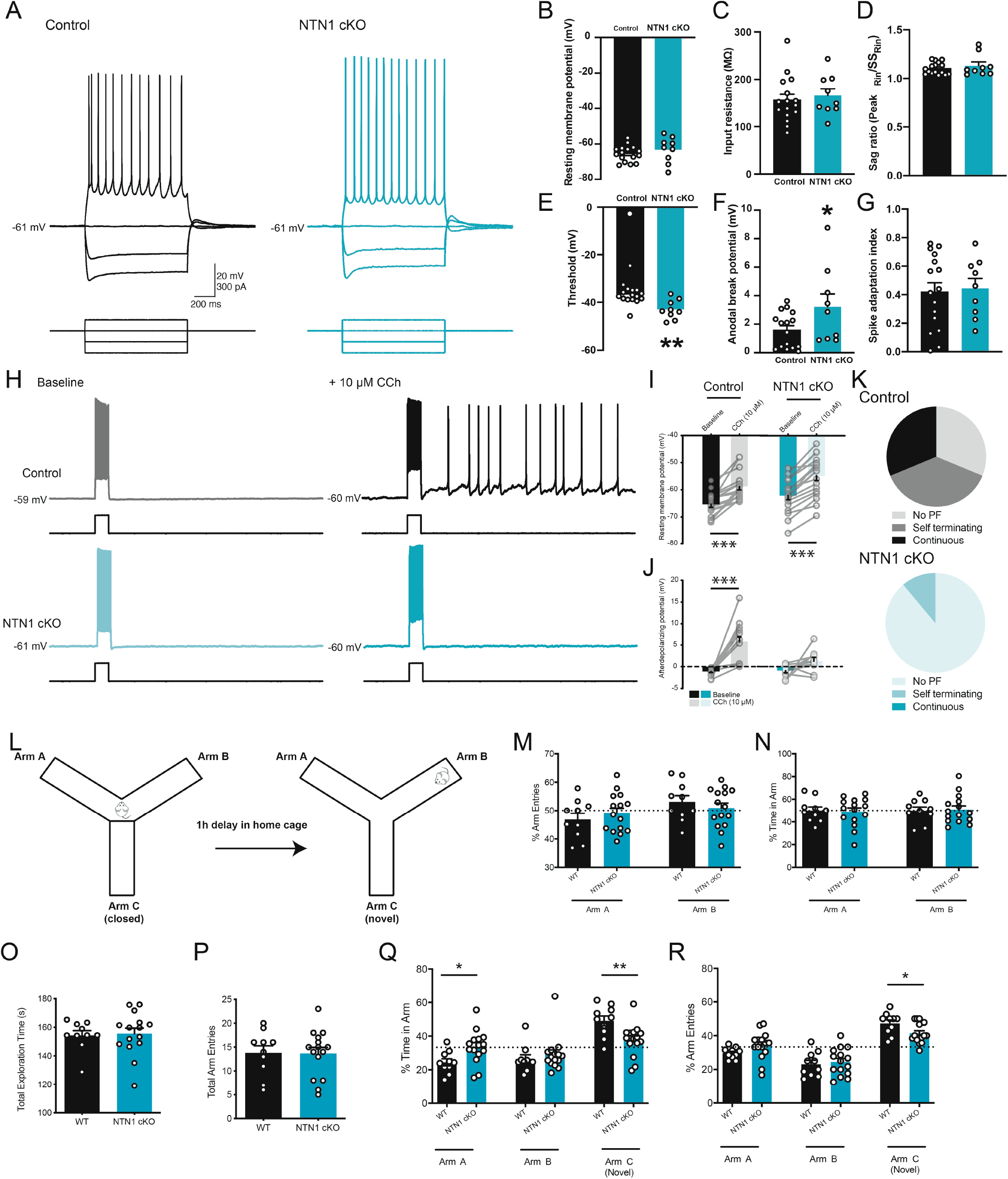
Cholinergic-mediated persistent firing activity is impaired in animals lacking netrin-1. A. Membrane voltage responses to hyperpolarizing and depolarizing intracellular current injections in layer V entorhinal neurons from control (black) and NTN1 cKO (blue) mice. B-G. Group data show no difference between control (black) and NTN1 cKO mice (blue) in resting membrane potential (B), input resistance (C), or sag ratio (D), but do show significantly reduced spike threshold (E; control: −36.6±1.1 mV, NTN1 cKO: −43.01±1.3 mV) and enhanced anodal break potential (F; control: 1.59±0.31 mV, NTN1 cKO: 3.22±0.90) with no change in spike frequency adaptation (G). H-K. Membrane potential response to a +100 pA intracellular current step (1 s) prior to (left) and following (right) bath application of CCh (10 μM) in a layer V entorhinal neuron from a control (black, top) and NTN1 cKO (blue, bottom) animal (H). Group data show that high concentrations of CCh (10 μM) significantly depolarize resting membrane potential (I; WT: −65.4±1.1 mV in baseline to −58.8±1.3 mV in CCh; NTN1 cKO: −62.1±1.5 mV in baseline to −54.8±1.7 mV in CCh). Group data also show that CCh significantly increases the amplitude of sADP in layer V entorhinal neurons from wildtype mice but not NTN1 cKO mice (J; WT: −1.12±0.21 mV in baseline to 5.81±1.09 mV in CCh; NTN1 cKO: −0.94±0.50 mV in baseline to 1.23±0.88 mV). Further, layer V neurons from wildtype mice showed persistent firing (11/16 cells, 68.7%), in contrast to only 11.1% of cells from NTN1 cKO (2/11, 11.1%) (K). L. Schematic diagram of the modified Y-maze test. M-N. Control (black) and NTN1 cKO (blue) mice show equal number of arm entries (M) and time spent in available arms (N) during baseline phase. O-R. Control (black) and NTN1 cKO (blue) mice show equal amount of time (O) and number of arm entries (P) during test phase. However, NTN1 cKO spend significantly less time exploring (Q) and make fewer entries into (R) the novel arm, *C*, compared to previously explored arms *A* and *B*. * denotes *p*<0.05, ** denotes *p*<0.01, *** denotes *p*<0.001.

Persistent firing has been hypothesized to underlie working memory, and has been linked to spatial navigation through the activity of grid cells in the entorhinal cortex (Hasselmo and Brandon, 2008). We have previously shown that netrin-1 is necessary for long-term spatial memory retrieval, but it remained unclear whether netrin-1 may also contribute to short-term memory storage (Wong et al., 2019). To assess the role of netrin-1 in short-term working memory, we used a novel modified non-food contingent delayed-nonmatch-to-sample Y-maze that exploits intrinsic novelty-seeking behavior of mice (Fig. 4L). During the sample phase, mice were allowed to freely explore two arms of the maze for 5 min. Both control and NTN1 cKO mice made similar numbers of arm entries and spent equal time in each arm (Fig. 4M-N). Animals were returned to their home cage for 1h, after which they were returned to the Y-maze and allowed to explore all arms of the apparatus, including the previously-closed arm. Control animals made significantly more entries and spent significantly more time in the novel arm compared to the familiar arms, due to their preference for novelty (Pogorelov et al., 2005). In contrast, NTN1 cKO mice did not show any arm entry preference and spent equal time in all three arms (Fig. 4O-R). These findings suggest that conditional genetic deletion of netrin-1 from forebrain principal neurons results in impaired spatial working memory.

Levels of acetylcholine are elevated during active spatial exploration in rodents (Marrosu et al., 1995). Interestingly, sustained cellular activity has been postulated to play a critical role in delayed non-match-to-sample tests and associated working memory functions such as spatial periodicity underlying place cell and/or grid cell ensemble formation (Hasselmo, 2008). Behavioral time-scale synaptic plasticity (BTSP) elicits bistable up- or down-states in CA1 pyramidal neurons, which may regulate *de novo* spatial representations through the emergence of place or grid cell activity (Bittner et al., 2017). It is possible that the relative genetic homogeneity of deep layer entorhinal neurons may facilitate plasticity events associated with spatial representations however it remains unclear how these factors may contribute to overall network activity (Ramsden et al., 2015).

Both netrin-1 and DCC are expressed throughout the deep layers of the entorhinal cortex, and have been previously demonstrated to play critical roles in the organization of hippocampal-entorhinal interactions (Barallobre et al., 2000; Lein et al., 2007). Moreover, they are critically implicated in N-methyl-D-aspartate receptor (NMDAR)-dependent forms of synaptic plasticity and spatial memory formation, and netrin-1 release can trigger rapid recruitment of Ca^2+^-permeable AMPARs to strengthen nascent synapses (Glasgow et al., 2018; Horn et al., 2013; Wong et al., 2019). Cholinergic recruitment of plasma membrane DCC sensitizes neurons to activity-dependent release of netrin-1, which can lead to the rapid modification of synapse strength that underlies cellular spatial representations. In contrast, selective loss of cholinergic tone through deafferentation is likely to weaken the propensity of synapses to undergo plastic changes (Schmitz et al., 2016). This raises the possibility that the slow actions of netrin-1 are necessary for more rapid cholinergic plasticity that is required for synaptic maintenance.

Early stages of Alzheimer’s disease (AD) are characterized by deafferentation of cholinergic fibers in the entorhinal cortex and hippocampus originating from the basal forebrain, which has led to the cholinergic hypothesis of AD (Francis et al., 1999; Schmitz et al., 2016). Consistent with this idea, optogenetic stimulation of cholinergic fibers can enhance synaptic plasticity in the hippocampus in a timing-dependent manner (Gu and Yakel, 2011). Our findings indicate that acetylcholine promotes DCC plasma membrane recruitment, and that the synergistic actions of acetylcholine and netrin-1 drive synaptic plasticity underlying both short- and long-term learning and memory processes.

## Acknowledgments

The authors would like to thank Nathalie Marcal, Kurt Soderstrom, and Mahmoud Moussa for assistance with genotyping, as well as all members of the Kennedy, Ruthazer, and Séguéla labs for their helpful comments on the manuscript.

## Author contributions

Conceptualization, S.D.G., J.G., P.S., E.S.R., and T.E.K.; Methodology, S.D.G.; Investigation, S.D.G., E.W.W., I.V.B., K.L., and J.G.; Formal Analysis, S.D.G., E.W.W., I.V.B., and K.L.; Writing – Original Draft, S.D.G.; Writing – Review & Editing, S.D.G., J.G., P.S., E.S.R., and T.E.K.; Funding Acquisition, P.S., E.S.R., and T.E.K.; Resources, S.D.G.; Supervision, P.S., E.S.R., and T.E.K.

## Data and materials availability

All data are available in the main text or the supplementary materials, and available from corresponding author upon reasonable request.

## Materials and Methods

### Animals

The study was conducted in accordance with the guidelines of the Canadian Council for Animal Care and approved by the McGill University Animal Care Committee. All animals were housed in group housing, and provided *ad libitum* access to food and water. For experiments using wild-type animals, C57/B6 mice (8-12 weeks) were used (Charles River, St Constant, Canada). Floxed alleles of *ntn1* (NTN1^*fl/fl*^ RRID:MGI:5755388) and *dcc* (DCC^*fl/fl*^) were maintained on a C57BL/6 genetic background as described (Bin et al., 2015; Horn et al., 2013). Animals that were homozygous for floxed alleles were crossed with T29-1 CaMKIIα-Cre mice (JAX 005359, RRID:IMSR_JAX:005359) from The Jackson Laboratory (Bar Harbor, ME, USA) and backcrossed for at least 5 generations on a C57BL/6 background. Both male and female T-29-CaMKIIα-Cre/netrin-1^*fl/fl*^ (T-29-CaMKIIα-Cre/NTN1^*fl/fl*^; NTN1 cKO, > 3 months old) and T-29-CaMKIIα-Cre/DCC^*fl/fl*^ (T-29-CaMKIIα-Cre/DCC^*fl/fl*^; DCC cKO, > 6 months old) mice were used for molecular biological and electrophysiology experiments. We observed no statistically significant differences between sexes, and therefore all data were pooled for analysis. Control experiments were performed using littermates that were negative for *Cre* and homozygous for floxed alleles of *ntn1* and *dcc*. For calcium imaging experiments, *dcc* null embryos were used.

### Hippocampal neuronal culture

For cell biological assays, cortices from E16.5 CD1 mice (Charles River, St-Constant, QC, Canada) were isolated, dissociated, plated and cultured as described (Goldman et al., 2013; Horn et al., 2013). Briefly, cortices were incubated at 37°C for 15 min in non‐supplemented Neurobasal (Invitrogen, Canada) containing trypsin (0.25%, Invitrogen, Canada). Tissue was triturated three times sequentially to yield a suspension of single cells. Cells were then plated in 6-well plates coated with poly-d-lysine (PDL) (20 μg/ml) and cultured for 7 DIV at a density of 1.5-2^^5^/mm^2^ in Neurobasal media containing: 1X B27, 1X N2, 0.4 mM Glutamine, 1 unit/mL penicillin, and 1 μg/mL streptomycin.

### Surface biotinylation

Biotinylation of cell surface proteins was performed as described previously (Bouchard et al., 2008). Cortical neurons from E16.5 mice were plated and cultured for 7 DIV at a density of 1.5-2 million cells per 10 cm PDL‐coated tissue culture dish. Briefly, cells were treated with carbachol (CCh) (0.1-100 μM, Sigma-Aldrich, cat no. C4382) for 15 min prior to biotinylation protocol. To test for dependence on muscarinic cholinergic receptors, cells were pre-treated with atropine sulphate (1 μM, Sigma-Aldrich, cat no. A0257) for 30 min prior to exposure to CCh. Cells were then washed twice with ice cold PBS (4° C) and subsequently transferred to cold PBS containing Ez-link Sulfo-NHS-LC-biotin (1 mg/ml; ThermoFisher Scientific) for 45 min. The biotin reaction was quenched by washing the cells twice for 25 min at 4°C in PBS containing 10 mM glycine, followed by 3x washes with cold PBS. The cells were then harvested in RIPA buffer containing 150 mM NaCl, 20 mM Tris, pH 8.0, 1%, NP-40 (USB Corp., Cleveland, OH, USA), 0.5% sodium deoxycholate, 0.1% SDS, 1 mM EDTA supplemented with protease inhibitors (1 mg/ml aprotonin, 1 mg/ml leupeptin, 100 mM PMSF, and 0.5M EDTA). Biotinylated proteins were precipitated using streptavidin-agarose beads (ThermoFisher Scientific) at 4°C for 2h, centrifuged at 13,800 x g for 15 min, and washed twice with cold PBS. Proteins were then eluted from beads using Laemmli sample buffer, and boiled for 5 min.

### Western Blots

Equal protein levels were loaded on acrylamide gels. Proteins were separated using SDS-PAGE on 8% polyacrylamide gels, and subsequently electroblotted onto nitrocellulose membranes. Non-specific binding was blocked in 5% non-fat milk diluted in PBS for 1h and subsequently incubated with primary antibodies overnight at 4°C. The primary antibodies used were: rabbit α-GAPDH (1:1000; Santa Cruz Biotechnology, RRID: AB_10167668), and mouse anti-DCC (1:1000; BD PharMingen, RRID:AB_395314). Blots were transferred to PBS containing horseradish peroxidase (HRP) conjugated secondary antibody (1:10 000) for 1h, and then developed and visualized by reaction with chemiluminesence reagent (Clarity Western Blotting Substrates Kit, BioRad) on radiography film (Carestream Blue X-ray film). Quantification of relative protein levels was performed on scanned images of immunoblots using Fiji (Schindelin et al., 2012).

### Calcium imaging

Calcium imaging of mouse hippocampal neurons was performed as described (Gibon et al., 2015). Briefly, cells were plated on poly-D-lysine coated 35 mm glass bottom dishes (MatTek) at a density of 100,000 cells / dish and incubated at 37° C in 5% CO_2_ until recording. After 7DIV, neurons were washed with Ringer solution containing: 140mM NaCl, 5mM KCl, 2mM CaCl, 1mM MgCl, 10mM N-2-hydroxyethylpiperazine-N’-2-ethanesulfonic acid (HEPES), 10mM glucose, (pH: 7.4), and loaded with Fura-2AM (5 μM) for 20 min. Cells were subsequently washed with Ringer solution for 30 min at 37° C. Neurons were imaged using an inverted TE2000-U microscope (Nikon) with a custom-built stage and equipped with a 40x oil-immersion objective (CFI super-S fluor, Nikon), and continuously perfused with Ringer solution throughout the entire recording. [Ca^2+^]_i_ signals were acquired using MetaFluor 7 software (Molecular Devices). All recordings were performed at rt, and various pharmacological antagonists were bath applied for a period of 10 min prior to baseline recording. Excitation light has been known to alter Fura-2 properties (Becker and Fay, 1987), and therefore all data were calibrated offline by subtracting control signal. Further, baseline measurements were obtained for >3 min prior to bath application of netrin-1 (2.7 nM), and all data are expressed as a function of baseline fluorescence ratio.

### *Brain slice* in vitro *electrophysiology*

Acute transverse hippocampal brain slices were obtained from wildtype (8-12 week old), NTN1 cKO (3-4 month old), and DCC cKO (6-10 month old) mice and their age-matched control littermates (Cre-negative floxed mice) using techniques previously described in detail (Glasgow et al., 2018). Briefly, mice were anaesthetized by I.P. injection of a mixture of 2,2,2 – tribromoethyl alcohol and tert-amyl alcohol diluted at 2.5% in PBS. Mice were then transcardially perfused with cold choline chloride-based solution (4° C) containing (in mM): 110 choline-Cl, 1.25 NaH_2_PO_4_, 25 NaHCO_3_, 7 MgCl_2_, 0.5 CaCl_2_, 2.5 KCl, 7 glucose, 3 pyruvic acid, and 1.3 ascorbic acid, constantly bubbled with carbogen (O_2_ 95%, CO_2_ 5%). Whole brain was dissected into choline solution, and thick horizontal brain slices (250 μm) containing the hippocampus and entorhinal cortex were cut using a vibrating microtome (VT1000s, Leica). Individual brain slices were allowed to recover for 1h in regular artificial cerebrospinal fluid (ACSF) containing, in mM: 124 NaCl, 5 KCl, 1.25 NaH2PO4, 2 MgSO4, 26 NaHCO3, 2 CaCl, and 10 glucose bubbled with carbogen (pH ~7.3, 300 mOsm) at rt (22°-24° C) prior to recordings.

For electrophysiological recordings, individual slices were placed in a custom-built recording chamber, and perfused with warmed (30 ± 2° C) ACSF (TC324B, Warner Instruments). Current and voltage recordings of visually- and electrophysiologically-identified layer V entorhinal cortex pyramidal neurons (Hamam et al., 2000) were performed on an upright microscope (Nikon Eclipse or Scientifica SliceScope 2000) equipped with a micromanipulator (Sutter MP-225 or Scientifica Patchstar), a 40x or 60x water immersion objective (0.8 or 1.0 N.A., respectively), differential interference contrast optics, and coupled to a near-infrared charge-coupled device camera (NC70, MTI or SciCam, Scientifica). Borosilicate glass pipettes (Sutter Instruments) (tip resistance: 4–8MΩ) were prepared using a horizontal puller (P-97, Sutter Instruments), and filled with an intracellular solution containing (in mM): 120 potassium gluconate, 20 KCl, 10 N-2-hydroxyethylpiperazine N′-2-ethanesulfonic acid (HEPES), 7 phosphocreatine diTris, 2 MgCl_2_, 0.2 ethylene glyco-bis (β-aminoethyl ether) N,N,N′,N′tetraacetic acid (EGTA), 4 Na^2+^-ATP, 0.3 Na^+^-GTP (pH adjusted to 7.20–7.26 using KOH, 275–285 mOsm). Whole cell somatic recordings from layer V pyramidal neurons were typically performed in the presence of kyurenic acid (1-2 mM) and picrotoxin (100 μM) to block glutamatergic- and GABA_A_-ergic synaptic transmission, respectively. Whole-cell current and voltage recordings were obtained using an Axopatch 200B or 700B amplifier (Molecular Devices). Current-clamp recordings were sampled at 20 kHz and filtered at 10 kHz, and voltage-clamp recordings sampled at 10 kHz and filtered at 2 kHz. All data were acquired through an analog-to-digital converter (Digidata 1322A or 1550A, Molecular Devices) for storage on computer hard disk with pClamp software (v9.0 or v10.4, Molecular Devices).

Access resistance was estimated on-line through compensation of the fast whole-cell capacitive transient elicited by a −5 mV voltage step using built-in compensation circuitry, set at 40%. Current clamp recording series resistance was compensated at 95%. Access and series resistances were calculated throughout the duration of the recording, and recordings were removed if the resistance was >30 MΩ and changed >20 %.

Intrinsic electrophysiological characteristics of layer V entorhinal pyramidal neurons were assessed using a series of hyperpolarizing and depolarizing intracellular current pulses (−200 pA to +100 pA). Neuronal input resistance was calculated by measuring the steady-state voltage response to a 100-pA hyperpolarizing current step (1 s). Sag ratio was calculated by expressing the peak input resistance as a function of steady-state input resistance. The anodal break potential was assessed as the peak voltage amplitude following the offset of a −200 pA hyperpolarizing current pulse, resulting in overshoot of the membrane potential. Basal excitability and resting membrane potential were routinely measured throughout the recording. Spike frequency adaptation was assessed using 1000-ms +100 pA current steps, and the adaptation index (AI) was computed according the formula: AI = 1−*f_f_*/*f_i_*, where *f_f_* is the final frequency, measured by final interspike interval (ISI) at end of intracellular current pulse, and *f_i_* is the initial frequency, measured using the ISI between first two action potentials (Reboreda et al., 2007).

Voltage-clamp recordings at −70 mV were used to assess ionic conductances associated with netrin-1 depolarization by use of current-voltage relationships. Holding voltage was varied between −130 mV and +50 mV in 20 mV increments. Netrin-1 elicited currents were then computed offline using subtraction of control IV relationships from IV relationships following netrin-1 application (Glasgow and Chapman, 2013). Pharmacological manipulations were bath applied for >10 min prior to bath application of netrin-1. Liquid junction potential of ~−13 mV was not corrected.

Plateau potentials and persistent firing were induced using a 1-s, +100 pA current injection while the neuron was held at −60 mV. Firing frequency was calculated as described previously (Reboreda et al., 2007). Continuous firing was defined as sustained neuronal spiking and plateau potentials at >60 s post-stimulus.

Stocks were prepared at 1000x by dissolving tetrodotoxin citrate (T-550, Alomone labs), carbachol (c4382, Sigma Aldrich), 4-Ethylphenylamino-1,2-dimethyl-6-ethylaminopyrimidinium chloride (ZD7288; ab120102, Abcam), 1-[2-(4-Methoxyphenyl)-2-[3-(4-methoxyphenyl)propoxy]ethyl-1*H*-imidazole hydrochloride (SKF-96365; 1147, Tocris) in ddH_2_0, and picrotoxin (P1675, Sigma-Aldrich), flufenamic acid (4522, Tocris), nifedipine (1075, Tocris), and 2-Aminoethoxydiphenylborane (1224, Tocris) in DMSO. The final concentration of DMSO did not exceed 0.1%. All other salts and drugs were from Sigma-Alrich unless otherwise noted.

### Spontaneous Alternation Y-Maze

The spontaneous alternation Y-maze relies on a rodent’s attraction to explore novel environments and used to assess possible impairments in novelty-seeking exploratory behaviour (Sarnyai et al., 2000). Mice were placed in the center of the maze (arm dimensions: 30.5 cm L x 10.5 cm W x 20.5 cm H) and were free to explore all three arms for 8 mins. The arms were randomly assigned as: A, B, or C. The walls of arms were identical to ensure homogenous intramaze environment. To further reduce testing-related stress, a thin layer of bedding (~0.5 cm) covered the floor of the maze. During exploration phase, the number and sequence of arm entries was recorded, defined as entry of all four paws inside the arm and their nose oriented toward the end of the arm. A spontaneous alternation occurred when a mouse entered a different arm of the maze in 3 consecutive arm entries. Spontaneous alternation (%) was calculated with the following formula:

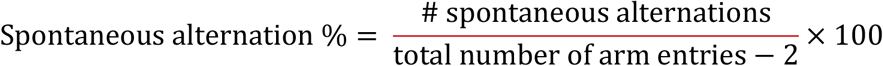

With a total of 11 arm entries, the percent spontaneous alternation would be approximately 56%.

### Modified Delayed Non-Match to Sample Y-Maze Task

The delayed non-match task (DNMT) is commonly used as a measure of working memory in rodents by relying on their natural tendency to alternate their goal choice arm in a Y-maze (Beracochea and Jaffard, 1995). We developed a non-invasive DNMT protocol that eliminates food deprivation and relies solely on innate exploratory motivation. During the “Sample Phase”, mice were initially placed in the center of the Y-maze (arm dimensions: 30.5 cm L x 10.5 cm W x 20.5 cm H), with one arm blocked by a guillotine door. Mice were allowed 5 mins to explore the two opened arms, which contained small cues on the walls. Following a 1-hr delay, during the “Choice Phase”, the door was removed from the novel arm. Mice were then placed in one of the two familiar arms and were allowed to explore all three open arms for 3 mins. An overhead camera recorded the mice in the Y-maze throughout training and testing, and exploration time and number of arm entries were measured by an experimenter blind to the genotypes.

### Statistical Analyses

Statistical analyses on parametric data were assessed using two-way repeated measures analysis of variance (ANOVA) followed by Tukey post-hoc test, one-way ANOVA followed by Tukey’s pairwise comparison test, and independent *t*-tests where appropriate. Analyses on nonparametric data were assessed using two-tailed Mann-Whitney test. Normality, homoscedasticity, and outlier tests were performed on all datasets. Data for Fig. 3K-L were compared to control conditions. Data were analyzed using Clampfit 10.3 (Axon Instruments), MATLAB (Mathworks), Fiji (Schindelin et al., 2012), Prism 7 (Graphpad), and Sigmaplot 11 (Systat). Plotted data were then formatted in Adobe Illustrator CS6 (Adobe Systems).

---

**Supplemental figure 1.**
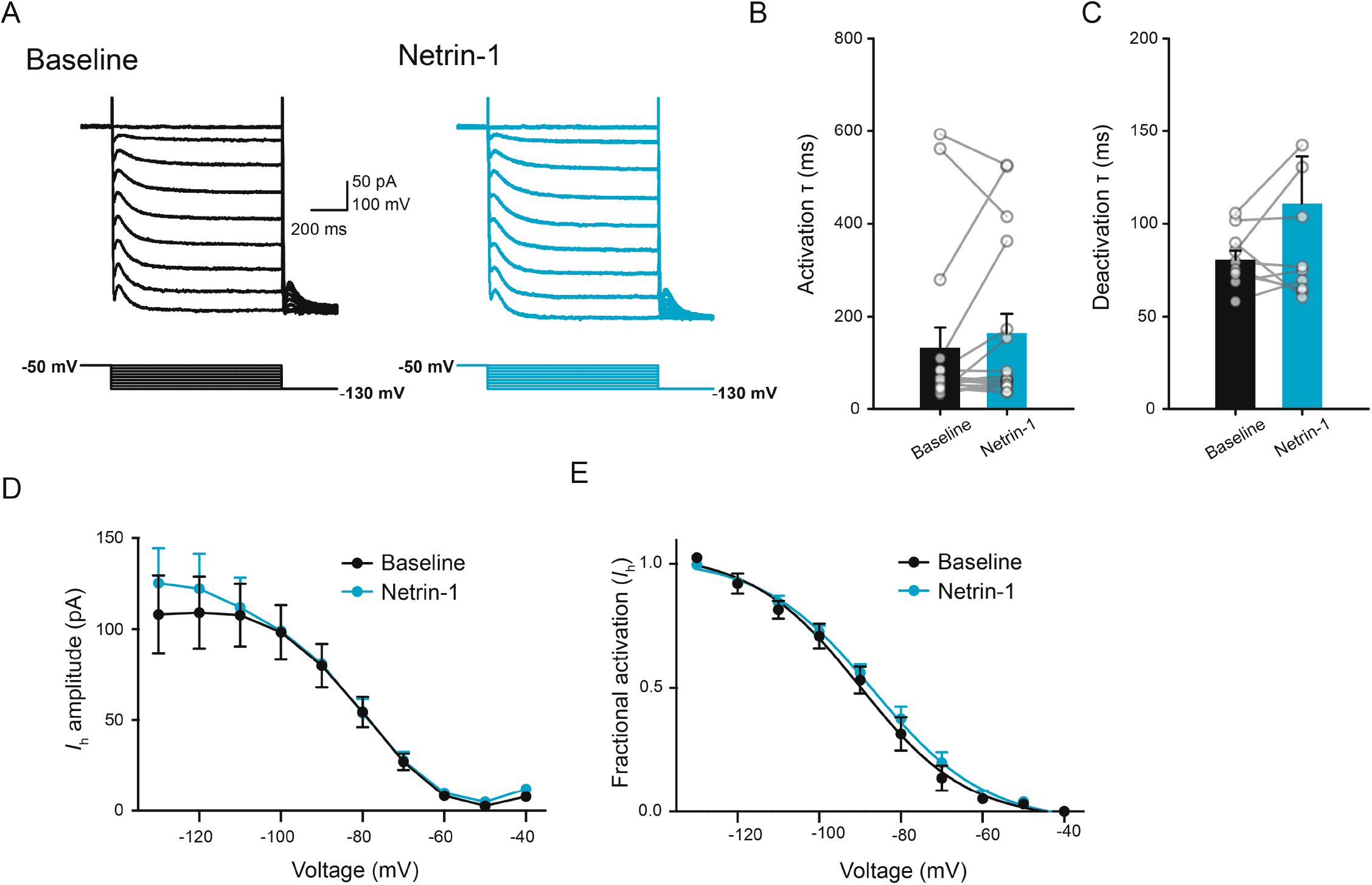
Netrin-1 does not impact *I*_h_. A. Current responses in a representative layer V entorhinal neuron prior to (black, left) and following bath application of netrin-1 (2.7 nM) (blue, right) across a range of voltages (−130 mV to −40 mV). B-C. Group data show no differences in activation (B) or deactivation (C) kinetics of *I*_h_ prior to (black) and following (blue) netrin-1. D-E. Group data showing current amplitude (D) and fractional activation (E) of *I*_h_ across a range of voltages prior to (black) and following bath application of netrin-1 (2.7 nM).

**Supplemental figure 2.**
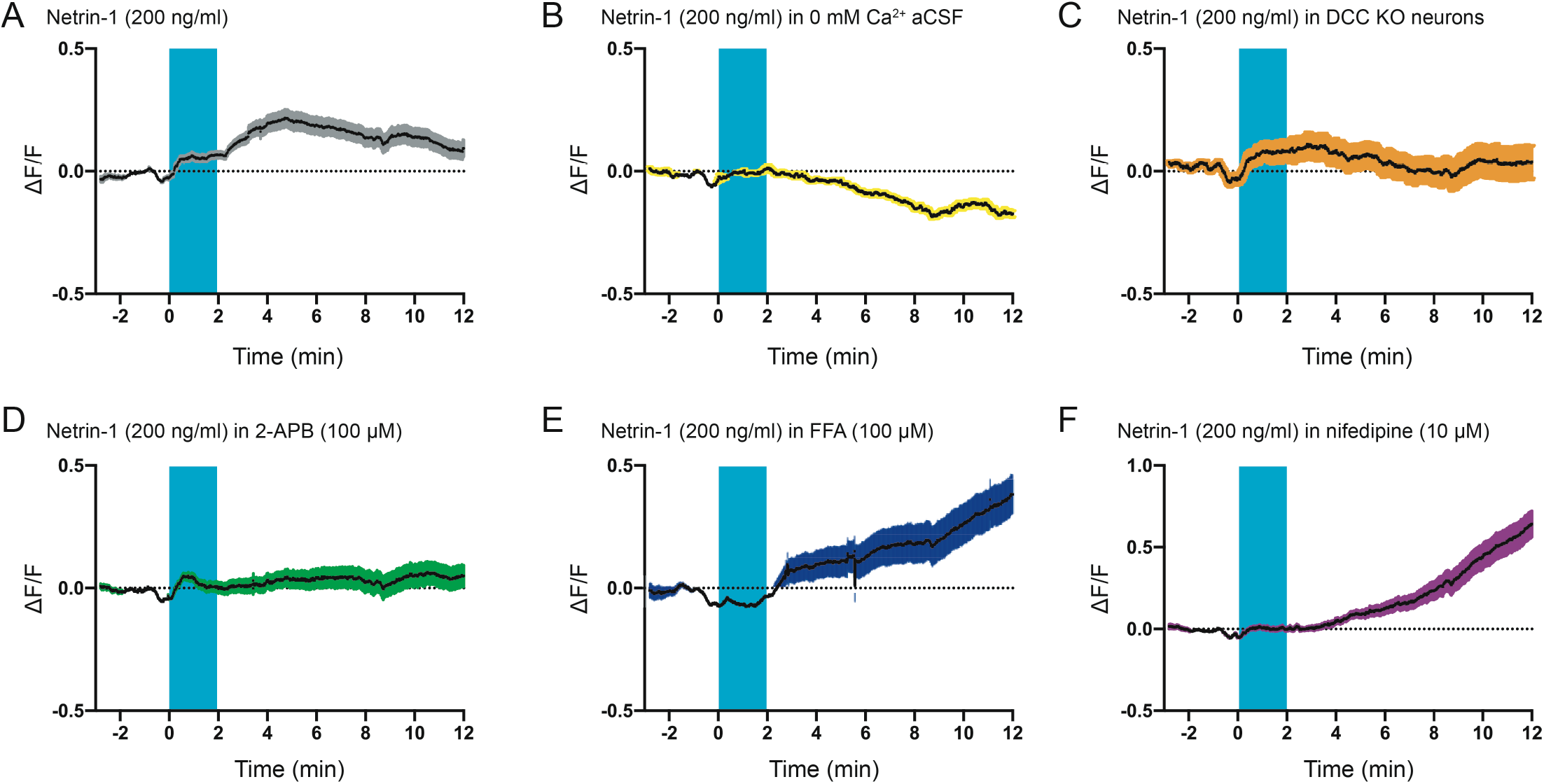
Netrin-1 induces DCC-dependent increases in [Ca2+]i in cortical neurons. A-F. Group data show that netrin-1 (2.7 nM) alone induces a long-lasting increase in fura-2 ratio (A) that requires extracellular Ca^2+^ (B), expression of netrin-1 receptor DCC (C), and is blocked by 2-APB (D; 100 μM), flufenamic acid (E; 100 μM), and nifedipine (F; 10 μM).

**Supplemental figure 3.**
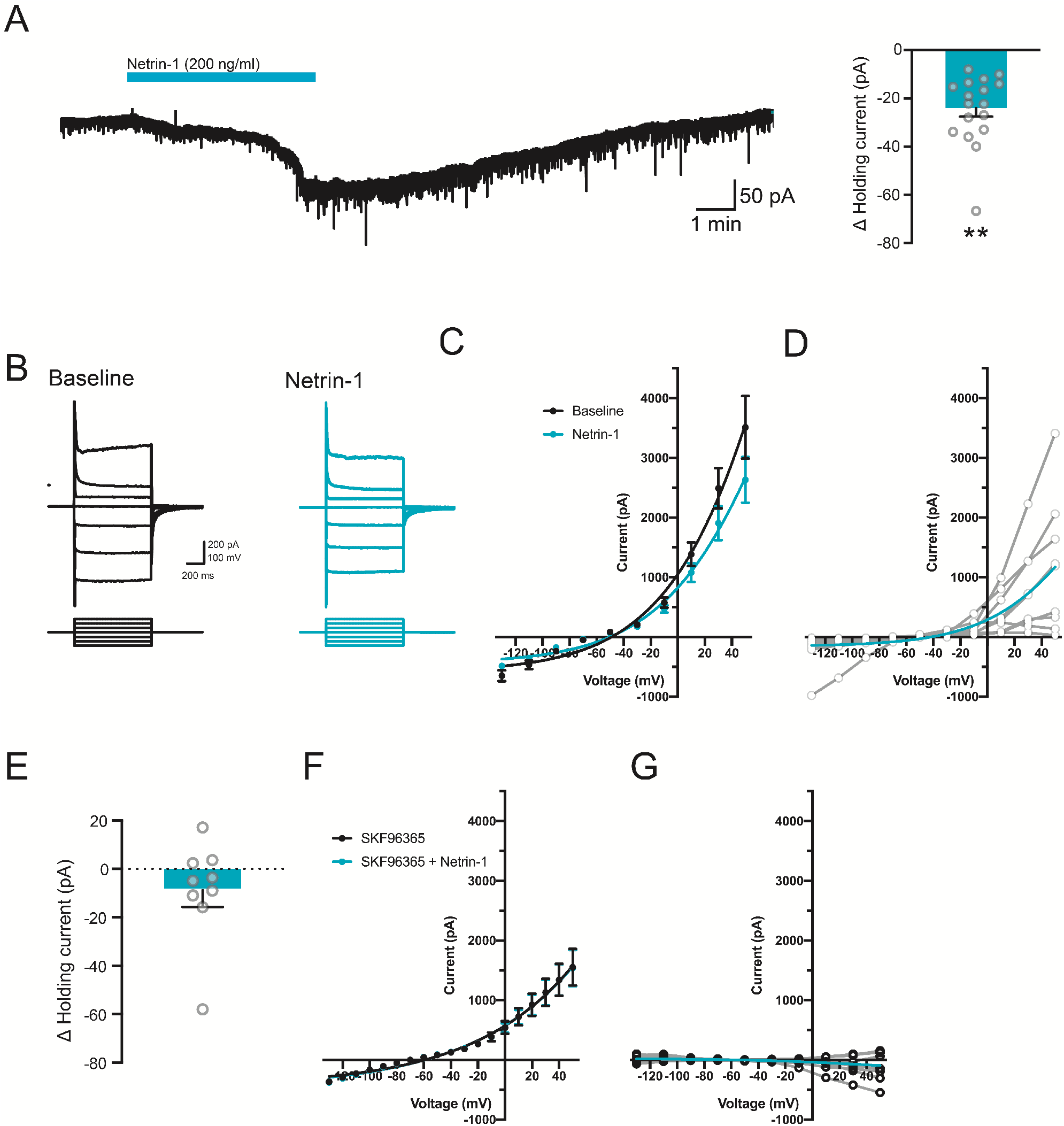
Netrin-1 induces a long-lasting inward current in layer V medial entorhinal cortex neurons. A. Representative current trace from a layer V entorhinal pyramidal neuron following bath application of netrin-1 (2.7 nM, blue bar) and group data show the change in holding current in layer V entorhinal pyramidal neurons held at −70 mV prior to and following netrin-1 (2.7 nM) (Δ current: −24.4±5.1 pA, right). B-D. Representative current-voltage traces (B), group data (C), and subtracted netrin-1 current (D) to a series of depolarizing and hyperpolarizing voltage steps in a layer V entorhinal neuron following netrin-1 (2.7 nM). Averaged (blue line) and individual (grey lines) subtracted current shown in D. E-G. Group data show the change in holding current (E; Δ current: −8.8±6.9 pA, *p* =0.24), current-voltage relationship to a series of depolarizing and hyperpolarizing voltage-steps (F), and subtracted current (G) following netrin-1 (2.7 nM) in the presence of TRPC blocker, SKF96365 (100 μM).

**Supplemental figure 4.**
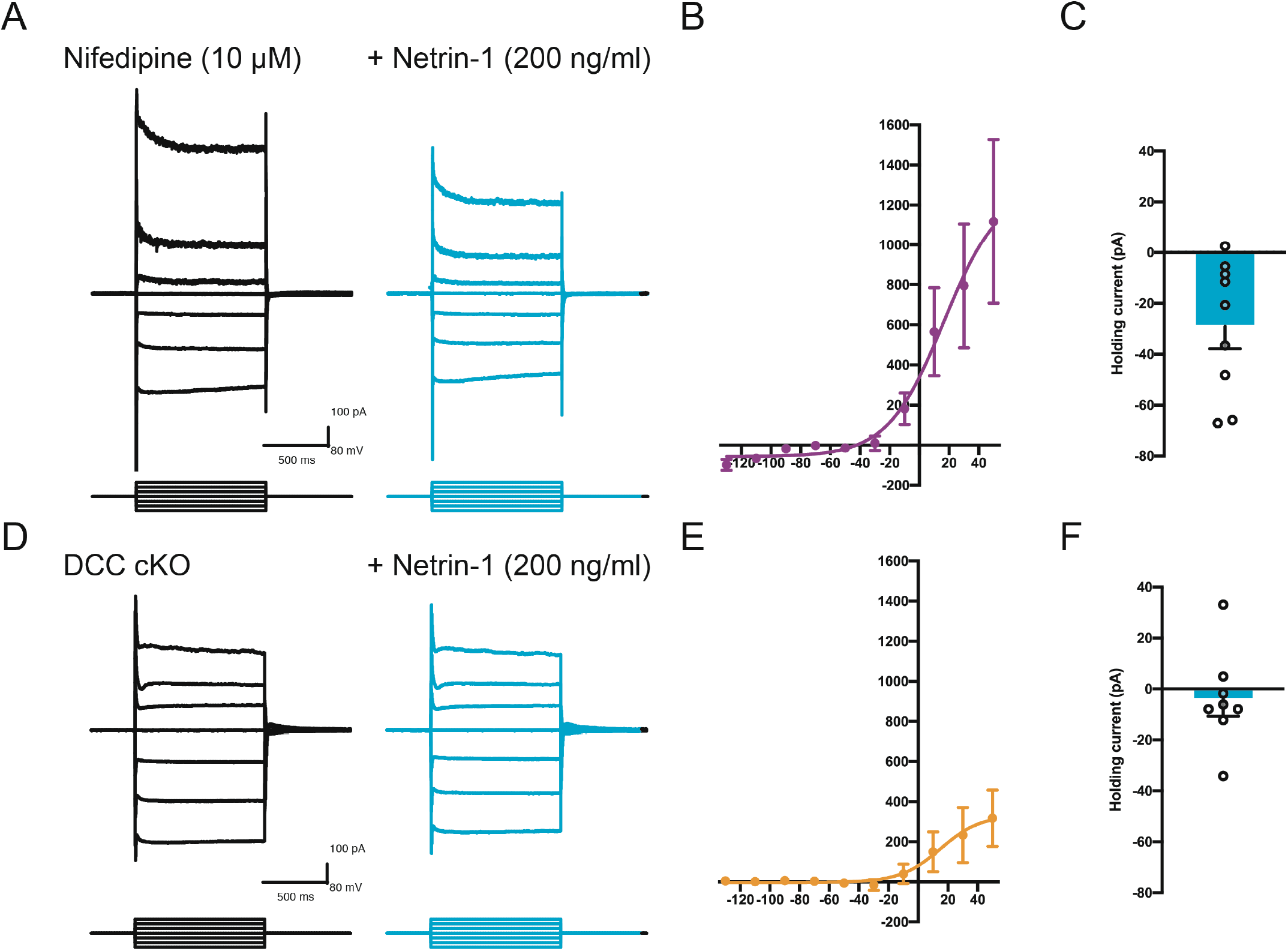
Netrin-1 induces an inward current that is dependent on netrin-1 receptor DCC but not L-type Ca^2+^ channels. Current responses in a layer V entorhinal neuron to a series of hyperpolarizing and depolarizing voltage steps prior to (black) and following bath application of netrin-1 (2.7 nM, blue) in the presence of nifedipine (A; 10 μM), and conditional deletion of DCC (D; DCC cKO). B, E. Group data show current subtraction of responses in the presence of netrin-1 from baseline values and holding current at −70 mV in the presence of nifedipine (B-C) or DCC cKO (E-F).

